# Shank2 expression identifies a subpopulation of glycinergic interneurons involved in nociception and altered in an autism mouse model

**DOI:** 10.1101/2020.05.24.112052

**Authors:** Florian olde Heuvel, Najwa Ouali Alami, Hanna Wilhelm, Dhruva Deshpande, Elmira Khatamsaz, Alberto Catanese, Sarah Woelfle, Michael Schön, Sanjay Jain, Stefanie Grabrucker, Albert C. Ludolph, Chiara Verpelli, Jens Michaelis, Tobias M. Boeckers, Francesco Roselli

## Abstract

Patients suffering from Autism Spectrum Disorders (ASD) experience disturbed nociception in form of either hyposensitivity to pain or hypersensitivity and allodynia. We have determined that Shank2-KO mice, which recapitulate the genetic and behavioural disturbances of ASD, display increased sensitivity to formalin pain and thermal, but not mechanical allodynia. We demonstrate that high levels of Shank2 expression identifies a subpopulation of neurons in murine and human dorsal spinal cord, composed mainly by glycinergic interneurons and that loss of Shank2 causes the decrease in NMDAR in excitatory synapses on these inhibitory interneurons. In fact, in the subacute phase of the formalin test, glycinergic interneurons are strongly activated in WT mice but not in Shank2-KO mice. As consequence, nociception projection neurons in lamina I are activated in larger numbers in Shank2-KO mice. Our findings prove that Shank2 expression identifies a new subset of inhibitory interneurons involved in reducing the transmission of nociceptive stimuli and whose unchecked activation is associated with pain hypersensitivity. Thus, we provide evidence that dysfunction of spinal cord pain processing circuits may underlie the nociceptive phenotypes in ASD patients and mouse models.

## Introduction

Autism Spectrum Disorders (ASD) are characterized by a range of sensory abnormalities, in particular in the domain of tactile sensitivity (Tomchek and Dunn, 2007). Among these, abnormal nociception is strikingly common in ASD and it manifests itself either as hypo-sensitivity or as hyper-sensitivity to painful stimuli. The common occurrence of self-injury and self-mutilation (including cases of self-extraction of teeth) and unreported wounds (Moore, 2015), together with clinical and experimental studies (Duerden et al., 2015; Klintwall et al., 2011; Tordjman et al., 2009) has been interpreted as evidence of reduced sensitivity to painful stimuli in ASD patients. Conversely, a subset of ASD patients display pain hypersensitivity, in the form of mechanical allodynia (Fründt et al., 2017) reduced threshold for thermal pain (Cascio et al., 2008) or pressure pain (Riquelme et al., 2016; Chen et al., 2017) and chronic pain unrelated to medical conditions (Clarke, 2015; Loades, 2015; Bursch et al., 2004). Pain hypersensitivity may constitute a major and underappreciated source of discomfort for ASD patients, in particular in those unable to properly report it because of reduced communication capabilities (Moore, 2015) and may lead to unnecessary medical procedures (Clarke, 2015).

The knowledge about the circuit mechanisms responsible for the altered nociception in ASD is limited. It is worth noting that ASD mouse models carrying mutations in different genes may display hyper- or hyposensitivity to pain, suggesting that the variability of the clinical phenotype may be linked to the genetic heterogeneity of ASD (more than 800 genes are linked to ASD; Vorstman et al., 2017). In fact, each genetic mutation might disrupt a discrete but different nodes of the nociceptive circuit, leading to mutation-specific phenotypes and mechanisms. In particular, loss of the ASD-related Shank3 protein results in pain hyposensitivity due to the disruption of the scaffold architecture enabling TRPV1 signaling in DRG neurons (Han et al., 2016).

Although substantial processing of proprioceptive and nociceptive stimuli takes place in circuits in the dorsal spinal cord (Braz et al., 2014), we largely ignore the extent of the involvement of these circuits in ASD-associated nociceptive phenotypes, either in terms of the cellular subpopulations involved or in terms of molecular and neurochemical abnormalities at work. One could speculate that, since a large fraction of ASD-associated genes code for synaptic proteins (Nelson et al., 2015), changes in the synaptic architecture, connectivity and excitation/inhibition balance may occur in the spinal cord of ASD patients and murine models.

Here we consider the ASD model obtained by deletion of the gene coding for the Post-Synaptic Density (PSD)-enriched scaffold protein Shank2 (Sheng and Kim, 2000). Indeed point mutations and missense mutations in Shank2 are responsible for a small but consistent fraction of ASD cases (Bourgeron, 2015).

Shank2-KO mice (obtained by targeting exon-6 and exon-7; Schmeisser et al., 2012; Won et al., 2012) display reduced social interaction, increased anxiety and compulsive grooming and are considered a *bona fide* mouse model of ASD. Recent work has suggested that Shank2-KO mice may display abnormal nociception due to alteration of spinal cord circuits (Yoon et al., 2017). Once again the neuronal subpopulations and the circuit architectural features responsible for such phenotype remain to be investigated.

Here we show that Shank2-KO mice display hypersensitivity to formalin-induced pain; notably, we have identified (in murine and human spinal cord) a subpopulation of glycinergic interneurons characterized by very high expression levels of Shank2. Loss of Shank2 results in the reduction in synaptic NMDAR and in the blunted recruitment of these inhibitory interneurons upon painful stimuli, which results in the over-activation of lamina-I nociceptive projection neurons.

## Materials and Methods

### Animals

All experiments were performed in agreement with the local and national guidelines for animal experimentation. In particular, all experiments were approved by the Regierungspräsidium Tübingen under the licence no o.103 and by the Italian Ministry of Health966/2016-PR.

All transgenic mouse lines used were previously described. GlyT2-EGFP mice tissue samples were a courtesy of Silvia Arber; spinal cord samples from PV-Cre x tdTomato-ROSA26 were a courtesy of Pico Caroni. Vgat-Cre x tdTomato-ROSA26, Ptf1a-Cre x tdTomato-ROSA26, Prrxl1-Cre x tdTomato-ROSA26 (also known as DRG11; Bechara et al., 2015) and vGluT2-CRE x tdTomato-ROSA26 double transgenic mice were a courtesy of Filippo Rijli, Ahmad Bechara and Alberto Loche. Ret-GFP transgenic mice were previously reported (Hoshi et al., 2012). Shank2-KO(Δ7) mice were previously described (Schmeisser et al., 2012).

### Shank2 antibodies

Polyclonal rabbit antiserum against the C-terminus of Shank2 (SA5192) was previously described (Boeckers et al., 1999). Briefly, partial cDNAs of the ProSAP1 cDNA (encoding amino acids 826–1259) were subcloned into the bacterial expression vector pGEX-1T (Pharmacia Biotech, Uppsala, Sweden). A 95 kDa glutathioneS-transferase (GST)-ProSAP1 fusion protein was expressed in Escherichia coli XL 1 Blue and used to immunize rabbits.

### Western blot

Western blot for Shank2 was performed as previously reported (Roselli et al., 2009). Briefly, lumbar spinal cord, cortex and cerebellum were dissected from WT and Shank2-KO mice (sacrificed by cervical dislocation). Tissue samples were quickly homogenized using a ?? in complete RIPA buffer additioned with Protease and Phosphatase inhibitors. The homogenate was subject to sonication and then cleared by centrifugation (10.000 g, 10 min, 4°C). Protein concentration was measured by BCA and 20μg of protein were loaded in each lane of an 8% -acrylamide gel. Proteins were transferred to a nitrocellulose membrane using a Trans-Blot^®^ Turbo^™^ Transfer System (Bio-Rad) semi-dry transfer apparatus (standard protocol for mixed molecular weight was run twice); membranes were blocked in TBS added with 1% Tween20 and 5% BSA for two hours and incubated with anti-Shank2 SA5192 antibody (diluted 1:500 in blocking solution) overnight at 4°C on an orbital shaker.

### Intraspinal injection of AAV9-GFP for sparse labelling

Intraspinal injection of AAV9 was performed as previously reported (Saxena et al., 2013). Briefly, WT mice aged P20 were administered buprenorphine (0.05mg/Kg) and meloxicam (0.1 mg/Kg) 30 min before being anesthetized with sevoflurane (4% in 96% O2, 800 ml/min). Dorsal skin was shaved and incised (10 mm) at lumbar level.

Paraspinal muscles were blunt-dissected and dorsal laminae were sectioned at vertebrae T11-T13 level. Upon removal of the dorsal laminae bone flap, the underlying dura was opened using a 33G needle and washed with ACSF. Viral injections were performed at the coordinates (y=+0.22-25mm; z=-0.55-6mm) having the central dorsal artery as reference. A total of four injections were performed, 250 nl/5 min each, using a pulled-glass capillary connected to a Picospritzer-III apparatus. Muscles were thereafter sutured on the midline using Prolene 7.0 surgical thread; the fascia was repositioned to cover the wound, and the skin was stiched on the midline using Prolene 6.0 surgical thread. Mice were then transferred to a warmed recovery cage with facilitated access to food and water and were administered additional doses of buprenorphine every 12h for the following 72h.

### Immunostaining

Spinal cord samples were processed as previously reported (Ouali-Alami et al., 2018). Briefly, mice were perfused with 4% PFA in PBS, (L1–L5) lumbar spinal cord was isolated and post-fixed for 18h in 4%PFA. Tissue samples were thereafter washed and cryoprotected overnight at 4°C in 30% sucrose. After embedding in OCT (Tissue-Tek), 40 μm cryostat sections were obtained. After blocking, primary antibodies (see supplementary table 1) were applied in PBS with 3% BSA, 0.3% Triton X-100, and incubated either 48-72h at 4°C on an orbital shaker. Sections were then washed with PBS (30 x 3 min at RT) and incubated for 2h at RT with appropriate combinations of secondary antibodies (see supplementary table 1) and DAPI (1:500) for nuclear staining (all from Invitrogen). After additional washing in PBS (30 x 3 min), the sections were dried and mounted in anti-fade Prolong mounting medium (Invitrogen). For STED Microscopy, before imaging, the samples were washed shortly with dH20 and mounted in 2,2-thiodiethanol (TDE) buffer.

### Immunostaining on Human spinal cord samples

Human spinal cord samples were obtained in agreement with the procedures approved by the ethical committee of Ulm University (n.245/17). and cryoprotected for 48h at 4°C in 30% sucrose. After Embedding in OCT (Tissue-Tek), 80 μm cryostat sections were obtained. Antigen retrieval was performed for 30 min at 100°C in citrate buffer, after which the sections were blocked (3% BSA, 0.3% Triton X-100 in PBS) and primary antibody incubation was performed for 72h at 4°C on an orbital shaker. Sections were washed in PBS (3 x 30 min at RT) and incubated for 2h at RT with the appropriate secondary antibody and DAPI for nuclear stain. After another round of washes (3 x 30 min at RT) the sections were dried and mounted in anti-fade Prolong mounting medium (invitrogen).

### In situ single-mRNA hybridization

Fluorescence in situ single mRNA hybridization (Wang et al., 2012) was performed according to manufacturer instructions (Fluorescence In Situ mRNA Hybridisation for fixed frozen tissue, RNAscope by ACDBio) with small modifications (as previously reported; olde Heuvel et al., 2019). Shortly, sections were mounted and dried on glass slides, quenching of autofluorescence was performed by pretreatment with 0.1M Glycine in PBS for 15 min. Thereafter, a 3 min antigen retrieval step was performed and sections were washed twice in dH2O and once in ethanol, followed by a pretreatment with pretreatment reagent III (all reagents were provided by ACDbio) for 20 min at 40°C. The Probes (GlyT2 and c-FOS) were hybridized for 4.5 h at 40°C, followed by a wash step for 2×2 min with wash buffer at RT. The sections were then incubated for 30 min with amplification-1 reagent at 40°C, followed by a wash step for 2×2min with wash buffer at RT. Whereafter the sections were incubated for 15 min with amplification-II reagent at 40°C, followed by a wash step of 2×2 min with wash buffer at RT. The last amplification step was performed by incubating the sections for 30 min with amplification-III reagent at 40°C, followed by a wash step of 2×2 min with wash buffer at RT. After this, the sections were incubated with amplification-IV for a final detection step for 45 min at 40°C, followed by a final wash step of 2×10 min with wash buffer. Sections were either counterstained with DAPI or a co-immunostaining was performed. For the co-immunostaining, the sections were blocked (10% BSA, 0.3% Triton in PBS) for 1 hour, followed by an overnight incubation at 4°C with primary antibody diluted in blocking buffer (see supplementary table 1). After the incubation the sections were washed for 3×30 min in PBS. Subsequently, a 2h incubation with secondary antibody diluted in blocking buffer at RT (Donkey anti-guinea pig 568, 1:500 [invitrogen]) was performed. After the last washing steps (3x 30 min in PBS), the sections were counterstained with DAPI and mounted using Fluorogold prolong antifade mounting medium (Invitrogen).

### Stimulated Emission Depletion microscopy (STED)

A custom build dual-color setup, with a high NA objective (HCX PL APO 100x/1.40-0.70 oil CS, Leica), was used for STED microscopy as described previously (Osseforth et al. 2014). Briefly, a super-continuum laser source (SC-450-PP-HE, Fianium) with a broad spectral range provided all the excitation (568 nm and 633 nm) and depletion (~720 nm and ~750 nm) beams, samples were scanned with a piezo stage (733.2DD, Physik Instrumente) and emission was recorded by an avalanche photodiode (SPCMAQRH-13/14-FC, Perkin-Elmer). Acquisition mode could be switched between confocal (diffraction limited resolution) or STED (lateral resolution ~35 nm) and was controlled by a custom LabVIEW (National Instruments) software.

### Confocal Imaging

Confocal images were acquired using a LSM Meta (Carl Zeiss AG) microscope as previously reported (Saxena et al., 2013), fitted with a 20× air objective or with 40x or 60x oil objectives with optical thickness fitted to the optimum value. For overview images, 20x objective 4×5 image tiles were acquired. For high-magnification images, a zoom factor 3x was applied during the acquisition of 60x oil objective images, oversampled in the z-axis to twice the theoretical optimal value. All images have been acquired at 1024×1024 pixels resolution and 12 bit depth. Acquisition parameters were set to avoid over or under-saturation and kept constant for each experimental set.

### Image analysis

For image analysis, confocal pictures were loaded into ImageJ. For the definition of Shank2 immunostaining intensity, a circular region of interest was manually located around each neuronal nucleus (identified in NeuN staining) and the integrated fluorescence intensity (expressed in arbitrary units -au-ranging between 0 and 4095) was logged. For the identification of Shank2^high^ neurons, we defined a threshold by considering the top of the distribution of intensities in laminae I and II and adding 200 au, thus defining as Shank2^high^ any neuron whose average fluorescence was at least 200 au higher than the brightest neuron in lamina I and II. Operationally, Shank2 immunofluorescence images were thresholded until neurons in lamina I and II were no longer visible and the threshold value was then moved 200 au higher; any Shank2+ neuron still visible at this stage was considered Shank2^high^. In the quantification of Shank2 expression in human samples, contour of neurons was manually drawn using the SMI321 immunostaining as reference and the average Shank2 immunostaining intensity was quantified. For the analysis of Homer, GluR2, vGluT1 and vGluT2 cluster size on GlyT2+ interneurons, the contour of each interneuron was manually drawn in imageJ; after thresholding, all clusters juxtaposed to the contour were highlighted and the size logged. The analysis of the c-Fos mRNA was performed by counting the single c-Fos mRNA dots per GlyT2+ interneurons (depicted with GlyT2 mRNA). Since c-Fos was expressed in every GlyT2+ neuron, we defined a threshold of at least 50 mRNA molecules per cell for it to be considered c-Fos positive. The neuronal and synaptic architecture was analyzed by taking a region of interest in the predefined lamina and counting manually the amount of neurons and with the Imaris software the density of synapses in the region of interest.

### Formalin test

Formalin test was performed according to the previously reported protocols (Wang et al., 2013; Lu et al., 2015). Briefly, animals were injected with 20μl 1% formalin solution in the dorsal surface of the left hindpaw; mice were thereafter swiftly put in a plexiglas chamber and video recording was obtained for 60 min and scored off-line for the duration of biting, licking or flinching activities. Phase I (0-15min) and phase II (20-45min) composite scores were computed.

### CFA-induced thermal hyperalgesia

Animals were injected with Complete Freund Adjuvant (100 μl, Sigma) subcutaneously into the left hindpaw, as described previously (lu et al., 2015). CFA injection led to an obvious tissue inflammation of the hindpaw characterised by erythema, oedema, and hyperpathia. For testing, each animal was placed in a box containing a smooth, temperature-controlled glass floor. The heat source was focused on a portion of the hindpaw, which was flush against the glass, and a radiant thermal stimulus was delivered to that site. The stimulus was shut off when the hindpaw moved, or after 20 s to prevent tissue damage. The time from the onset of radiant heat to the endpoint was the paw withdrawal latency (PWL). The radiant heat intensity was adjusted to obtain a basal PWL of 10–12 s. Thermal stimuli were delivered 3 times to each hindpaw at 5–6 min intervals.

### CFA-induced mechanical hyperalgesia

After CFA injection and induction of inflammation, von Frey filaments (VFFs) were employed to measure mechanical hyperalgesia according to established protocols (Chaplan et al., 1994). Briefly, a series of calibrated VFFs (0.4~25.0g) were applied perpendicularly to the plantar surface of the hindpaw with sufficient force to bend filaments for 60 s or until it withdrew. In the absence of a response, filament of next greater force was applied. In the presence of a response, filament of next lower force was applied. To avoid injury during experiments, cutoff strength of VFF was 25.0 g. The tactile stimulus producing a 50% likelihood of withdrawal was determined by means of the “up-down” calculating method, as described in detail previously (Chaplan et al., 1994). Each test was repeated 2 ~3 times at ~2 minute interval, and the average value was used as the force to induce a withdrawal response.

### Statistical Analysis

Statistical analysis was performed with the Graphpad Prism version 6.00 for Windows (GraphPad Software, La Jolla California USA) software package. Group comparisons were performed using the ANOVA test, with Dunnett’s or Bonferroni’s correction for multiple comparisons. When appropriate, non-parametric Mann-Whitney test with Tukey’s correction was used. All results are depicted as average±SD unless indicated.

## Results

### Shank2-KO mice display hypersensitivity to inflammatory pain and sensory modality-specific allodynia

First, we verified if Shank2-KO mice displayed abnormal nociceptive responses and pain processing using acute and injury-associated painful stimuli. We tested sensitivity to acute chemical/inflammatory nociception using the formalin test. The first phase of the formalin test pain response (between 0 and 10 minutes), a measure of nociceptors activation and transmission, was unaffected by Shank2 deletion: the cumulative time spent licking or flinching the injected hindpaw was comparable with the one in WT littermates (p>0.05; Figure 1A-B). Conversely, licking/flinching time in the phase II, related to short term spinal plasticity, was significantly longer in Shank2-KO mice (p<0.01; Figure 1A-B), indicating an increased pain perception (Lu et al., 2015; Foster et al., 2015).

**Figure 1:**
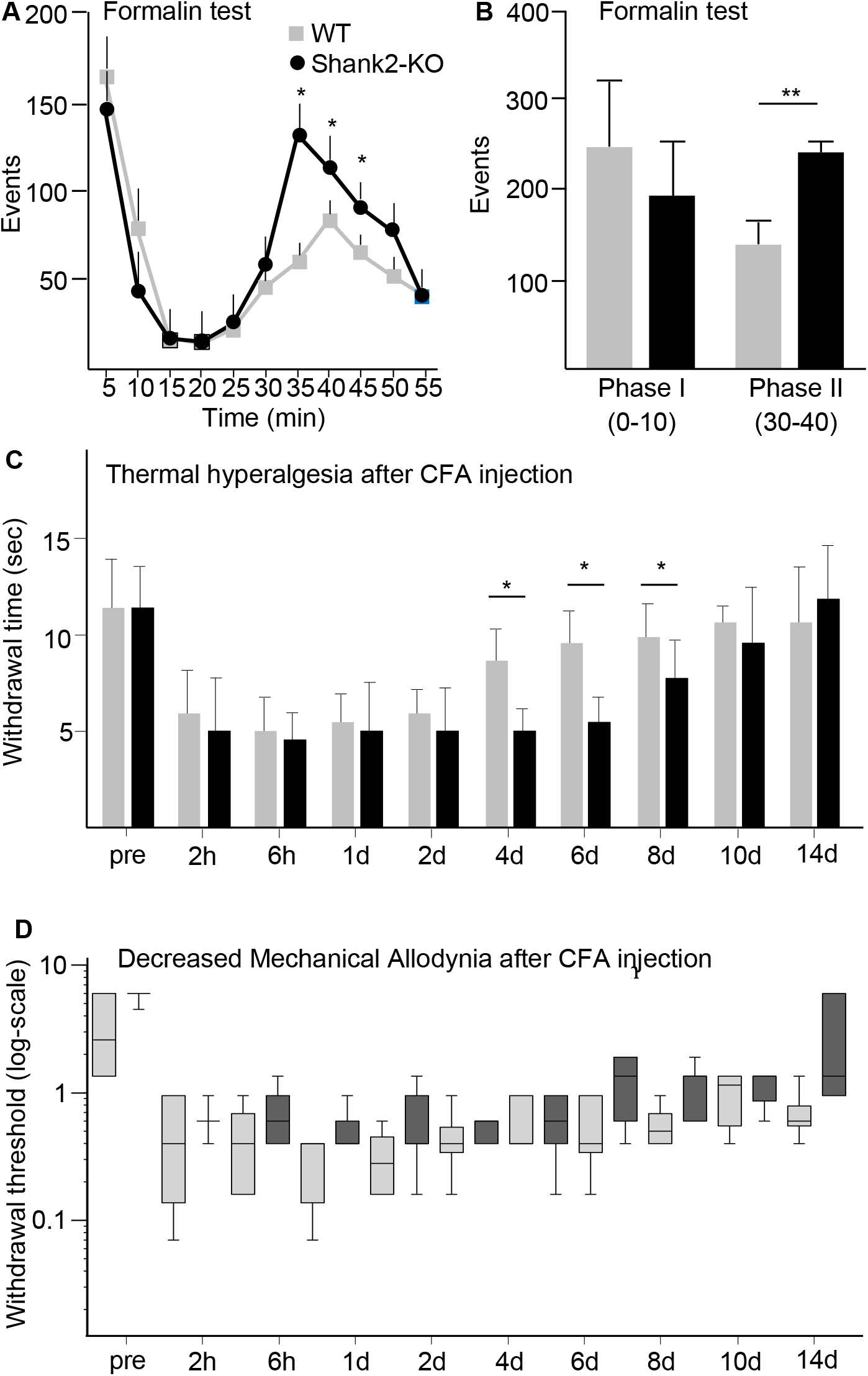
Shank2-KO mice display hypersensitivity to inflammatory pain and sensory modality-specific allodynia. Behavioural analysis of nociceptive responses in Shank2-KO and WT mice. A-B) Formalin was injected in the hindpaw to measure the licking/flinching response. Shank2-KO mice show no difference in licking/flinching in the first phase compared to WT (0-10min; 194±63 vs 243±41 respectively, p>0.05) after the formalin test. However, in the second phase (30-40min) a significant increase was observed in licking/flinching in the Shank2-KO mice (245±18 vs 141±127; KO vs WT; p<0.01). C) inflammation-induced thermal hyperalgesia was measured by Paw withdrawal time after CFA injection and subjection to a thermal source. Thermal hyperalgesia was found in both WT and Shank2-KO mice the first few days (1-4 days), normalisation of withdrawal time occured in the WT mice from 4 days, but a significant increased sensitivity remained in Shank2-KO mice until day 8 (day 4: 8.76±1.2 vs 5.07±0.83; day 6: 9.69±1.08 vs 5.51±0.72; day8: 10.02±0.68 vs 7.31±0.98; WT vs KO; p<0.05). D) Inflammation-induced mechanical hyperalgesia was measured by withdrawal threshold after CFA injection and von Frey filaments. Both WT and Shank2-KO mice show an increased sensitivity in the first few hours and days of the von Frey filaments. Shank2-KO mice show a trend toward decreased sensitivity at 6 days compared to WT mice. Data shown as average±SD, n=3.

We further tested thermal and mechanical hyperalgesia after chronic induction of inflammation by intraplantar injection of complete Freund’s adjuvant (CFA; Lu et al., 2015). After CFA injection, both WT and Shank2-KO mice displayed a comparable decrease in the latency of paw withdrawal from a thermal source, indicating the appearance of inflammation-induced thermal hyperalgesia; however, starting from day 4, WT mice displayed a rapidly progressive normalization of the withdraw latency, whereas Shank2-KO mice continued to manifest an increased sensitivity to thermal stimuli up to 8-10 days after CFA injection (WT vs Shank2-KO day 4: p<0.05; day 6: p<0.05; day 8 p<0.05; Figure 1C). Notably, thermal sensitivity before CFA administration and in the early phases of the inflammatory response was comparable in Shank2-KO and WT. Thus, only the recovery of the inflammation-induced thermal hyperalgesia was affected by the Shank2 deletion.

Conversely, baseline sensitivity to mechanical stimulation by von Frei’s filaments was indistinguishable in WT and Shank2-KO mice. After CFA injection, in contrast to thermal hyperalgesia and to the response to formalin injection, Shank2-KO mice did not display an increased sensitivity to mechanical stimulation and actually displayed a trend towards faster resolution of the inflammation-induced mechanical allodynia (thus, decreased mechanical allodynia; Figure 1D).

Taken together, these data show that Shank2-KO mice display a selective hypersensitivity to noxious chemical and thermal stimuli in the sub-acute/chronic phase but not in the acute phase, and that the hyperalgesia is sensory-modality-specific, since it appears for thermal stimulation but not for mechanical stimulation.

### A distinct Shank2^high^ neuronal subpopulation in dorsal and ventral spinal cord

In order to mechanistically investigate the origin of the abnormal nociception in Shank2-KO mice, we first analyzed the expression pattern of Shank2 in the spinal cord of WT animals. Immunostaining of spinal cord sections with the anti-Shank2 SA5192 antiserum revealed a punctuate pattern distributed throughout the gray matter, coherent with Shank2 synaptic localization. Notably, Shank2+ puncta appeared to delineate a subset of neurons scattered in dorsal laminae III, IV and V (as well as in the ventral horn; Figure 2A) characterized by dense and substantially more intense Shank2 immunoreactivity. We considered the distribution Shank2 immunostaining intensity of neurons in laminae I and II, which topped at 1000 au of intensity, and we introduced a 200 au margin and we defined all neurons with averaged Shank2 immunostaining intensity larger than 1200 as Shank2^high^ neurons (see Methods for the operational identification of Shank2^high^ neuron; Figure 2A-B). Supporting the immunolabelling result, expression data from the adult spinal cord in the Allen Spinal Cord Atlas (http://mousespinal.brain-map.org/imageseries/show.html?id=100026736) revealed a subpopulation of Shank2 high-expressing cells intermingled with other dorsal and ventral spinal cord neurons. The dorsal population of Shank2^high^ neurons could be further distinguished into two subgroups based on the medio-lateral localization of the neurons: one (smaller) located medially and one (larger) located more laterally within the laminae III, IV and V (Figure 2C-D).

**Figure 2:**
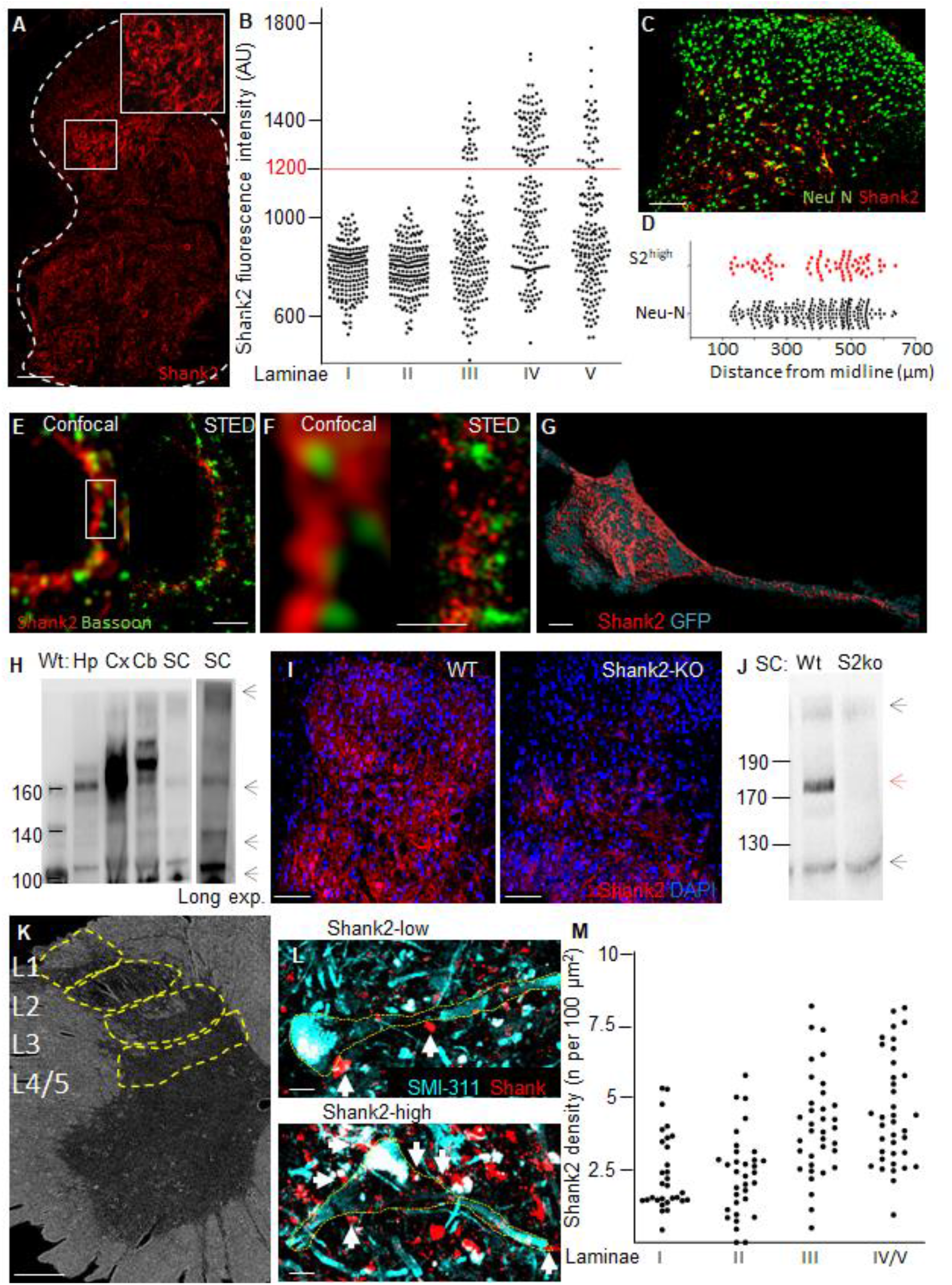
High Shank2 expression identifies a distinct neuronal subpopulation in spinal cord. A-B) Shank2 IF staining revealed a synaptic pattern around cells in the dorsal and ventral horn, with a subset of Shank2 high expression neurons in layer III, IV and V (mean intensity; 785±99, 788±100, 892±241, 1097±256, 941±235, Layer I-V respectively). C-D) These Shank2^high^ expressing neurons were further divided into location from midline in layer III, IV and V (median; 425±138 vs 375±122 Shank2^high^ vs Neu-N. E-F) High magnification confocal pictures reveals a continuous band of Shank2 around cells, however STED imaging shows the distinct punctuate patterns of Shank2 colocalised with Bassoon. G) sparse labeling of neurons in the dorsal lamina was performed by AAV9-CMV-GFP intraspinal injection, combined with Shank2 IF staining. Shank2 is highly distributed on cell-body and dendrites H) Shank2 Western blot reveals 4 isoforms in the spinal cord located at 220-240, 160, 140 and 100 kD. The isoforms 160 and 100 were also detectable in other CNS regions like cortex, hippocampus and cerebellum. The cerebellum also showed the 220-240 isoform which was not found in cortex or hippocampus. I-J) IF staining and western blot of WT and Shank2-KO mice show an almost complete loss of Shank2^high^ expressing neurons in layer III, IV and V, this loss corresponds to the loss of MW isoforms at 160 and 140 kD, the highest and lowest MW were still expressed. K-M) IF staining of Human spinal cord samples reveal expression of Shank2 across the cell body and dendrites, Lamina III and IV have neurons with a higher density of Shank2 expression on both cell body and dendrites similar to neurons in mouse spinal cord. Data shown as average±SD, n=3, Scalebar; A: 100μm, C: 50μm, E: 1μm, F: 500nm, G: 1μm, I: 100μm, K: 1000μm, L: 10μm.

Examined at high magnification, Shank2^high^ neurons showed Shank2 immunolabelling as an almost continuous band surrounding the cell body and proximal dendrites. Nevertheless, when samples immunostained for Shank2 (and synaptophysin) were imaged using Stimulated Emission Depletion microscopy (STED, Eggeling et al., 2013) at 60-nm-lateral resolution, the continuous Shank2 staining in Shank2^high^ neurons was resolved into a series of closely juxtaposed Shank2 clusters, of variable size, facing synaptophysin-positive presynaptic terminals (Supplementary Figure 1A-B). Furthermore, STED images revealed that on Shank2^high^ neurons each Shank2 cluster matched a single Bassoon-positive release site (Figure 2E-F and Supplementary Figure 1C-E). Thus, in Shank2^high^ neurons, Shank2 is distributed in high-density clusters involved in synaptic contacts.

In order to show the expression of Shank2 in Shank2^high^ neurons in compartments distinct from the cell body, we performed a sparse-labelling of dorsal spinal neurons by injecting in the spinal cord 200 nl of AAV9-CMV-GFP suspension; 15 days after injection we selected for analysis GFP-positive Shank2^high^ neurons. Confocal stacks were acquired at high magnification over the cell body and dendrites of the selected neurons, and reconstructed in 3D rendering (Figure 2G). Shank2 clusters were distributed also on dendrites, although at reduced density compared to the cell body, confirming that Shank2^high^ neurons have Shank2-enriched synapses across the whole neuronal structure.

The Shank2 transcript is known to undergo extensive alternative splicing, giving rise to multiple isoforms (Lim et al., 1999). In fact, spinal cord homogenates analyzed by WB (together with hippocampal, cerebellar and cortical samples) and probed with the SA5192 polyclonal antibody against the C-terminal domain (shared by all Shank2 isoforms; Boeckers et al., 1999) displayed one high-MW isoform (220-240 kDa; attributable to the ankyrin-repeats-containing 240 kDa shank2E isoform; also detectable in cerebellum but weak-to-indetectable in hippocampus and cortex) and three more isoforms at 160, 140 and 100 kDa, of which the first and the last were shared by cerebellum, cortex and hippocampus (Figure 2H). On the other hand, immunoblots of the spinal cord of Shank2-KO mice were characterized by the loss of the most abundant 160 and 140 kDa isoform (Figure 2J) but persistent expression of the low-abundance highest and lowest MW isoforms.

Immunostaining of spinal cord samples from Shank2-KO mice also revealed the almost complete loss in Shank2 immunoreactivity (Figure 2I), with limited residual immunopositive signal in the cell body of scattered neurons in laminae III, IV and V; this may be due to the residual Shank2 isoforms observed by WB. Immunostaining of post mortem human spinal cord revealed a shank2 expression in neurons, with a similar distribution of Shank2^high^ and shank2^low^ across various laminea compared to mouse (Figure 2K-M). Thus, Shank2 is expressed at substantial levels in the mouse and human spinal cord, and in particular in a previously unrecognized subpopulation of spinal cord neurons and it is no longer detectable in Shank2-KO mice.

### Shank2 ^high^ neurons are a subpopulation of glycinergic interneurons in dorsal laminae

In order to establish the neurochemical identity of the Shank2^high^ neurons, we immunostained for Shank2 an array of spinal cord samples from mice in which distinct spinal cord neuronal subpopulations have been genetically-labelled neuronal subpopulations: vGlut2-Cre;ROSA26-Tomato to mark excitatory neurons, GlyT2-GFP, VGAT-CRE;ROSA26-Tomato or PV-CRE;ROSA26-Tomato to label all or part of inhibitory interneurons. In laminae III and IV, only 2.4±0.8% of Shank2^high^ colocalized with Tomato+ neurons in vGlut2-Cre;ROSA26-Tomato mice, suggesting the inhibitory nature of Shank2^high^ cells (Figure 3B and 3H). In fact, 82.4±4.2% of Shank2 neurons were GFP+ in GlyT2-GFP mice (Figure 3A and 3H) and virtually all (96.6±1.7%) Shank2^high^ neurons was Tomato+ in VGAT-CRE;ROSA26-Tomato (Figure 3D and 3H). In PV-CRE/tdTomato mice, a small fraction (12.9±0.4%) of Shank2^high^ neurons corresponded to PV-positive interneurons (Figure 3C and 3H), almost all in the medial population (located in the *nucleus proprius* of the spinal cord). Nevertheless, Shank2^high^ cells represented only 71.1±4.1% of glycinergic interneurons (up to 30% of GlyT2-GFP cells and more than 80% of PV+ neurons were Shank2^low^, data not shown). We further characterized the transcriptional and developmental identity of Shank2^high^ neurons. In agreement with their inhibitory nature, Shank2^high^ neurons did not correspond to Prrxl1+ cells (excitatory interneurons involved in the processing of nociceptive inputs, Rebelo et al., 2010; Figure 3F and 3I) in Prrxl-Cre/TdTomato mice or to neurons originated from dorsal, Wnt1-positive progenitor populations (in Wnt1-cre;ROSA26-Tomato double tg mice; Hsu et al., 2010; Figure 3G and 3I). On the other hand, a significant fraction (66.0±12.3%; Figure 3E and 3I) of Shank2^high^ neurons correspond to Tomato+ cells in Ptf1a-Cre;ROSA26-Tomato mice; in fact, Ptf1a transcriptionally regulates the development of inhibitory interneurons phenotype and neurochemistry (Borromeo et al., 2014).

**Figure 3:**
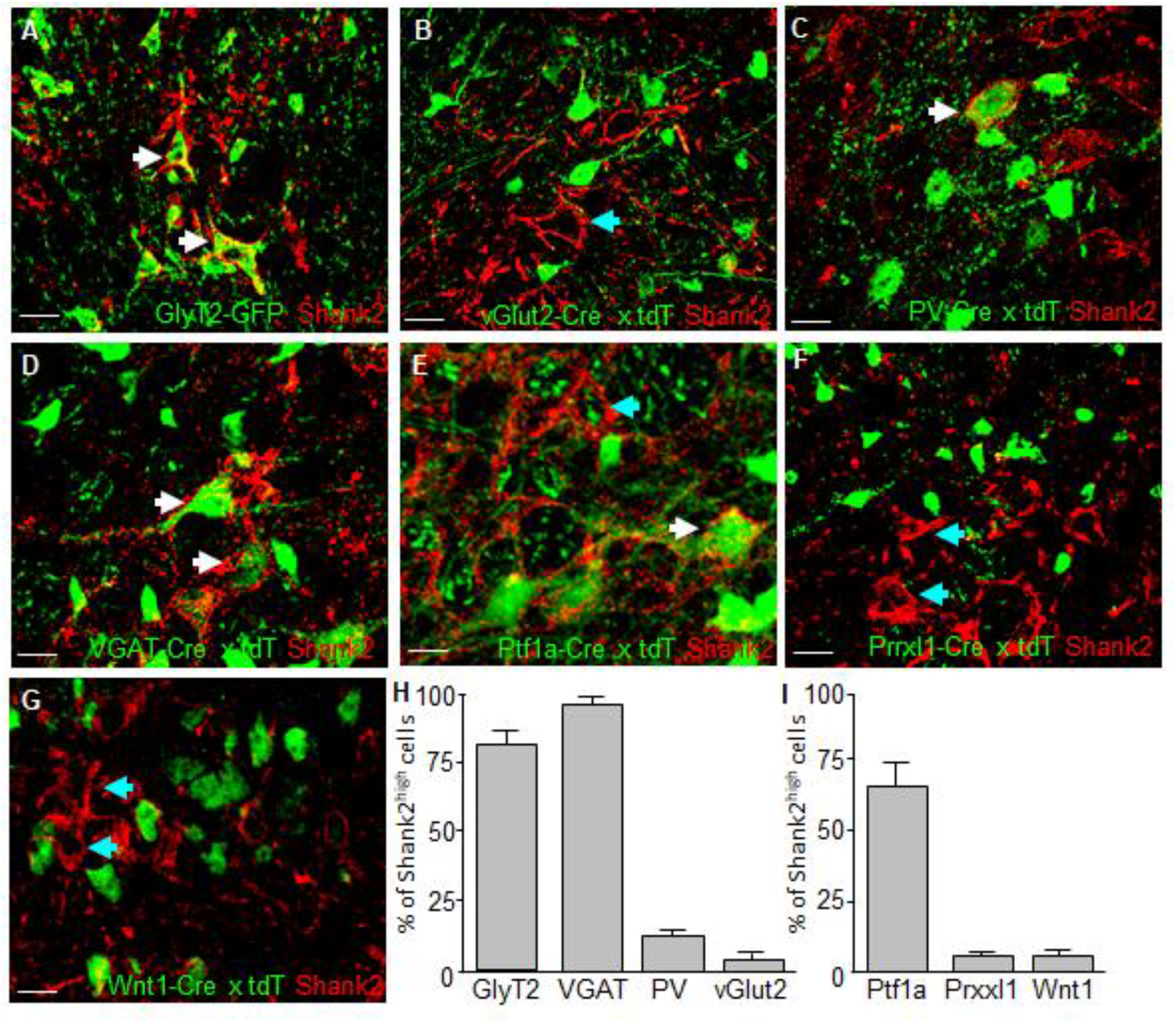
Shank2^high^ cells are mainly glycinergic interneurons. A subset of genetically labelled mice were used to investigate in which type of cell Shank2 was mainly expressed. AD and E) GlyT2-GFP, vGluT2-Cre;ROSA26-Tomato, VGAT-Cre;ROSA-Tomato and PV-Cre;ROSA26-Tomato mice were used to identify inhibitory and excitatory neurons in the dorsal horn of the spinal cord. Staining spinal cord sections with Shank2 revealed the inhibitory nature of Shank2 cells (96.6±0.4% vs 2.4±0.8% VGAT+ vs GlyT2+), with a high number of Shank2 expressing neurons on GlyT2+ cells (82,4±4,2%) and a lower amount of Shank2 expressing neurons on PV+ cells (12.9±0.4%). High expression of Shank2 was mainly observed in GlyT2+ cells compared to PV+ cells (70% vs 20%). E-G and I) Prrxl-Cre;Tomato, Wnt1-Cre;Tomato and ptf1a-Cre;Tomato mice were used to identify a subset of transcription factors regulating excitatory and inhibitory phenotypes. Staining spinal cord sections of these mice resulted in a low fraction of Shank2 expressing neurons on prrxl1+ and Wnt1+ cells and a high number of Shank2 expressing neurons on ptf1a+ cells (66±12.3%). Data shown as average±SD, n=3, scale bar: 20μm.

Taken together, these data prove that Shank2^high^ neurons represent a previously unrecognized subpopulation of inhibitory glycinergic/gabaergic interneurons in the sensory areas of the spinal cord.

**Supplementary Figure 1:**
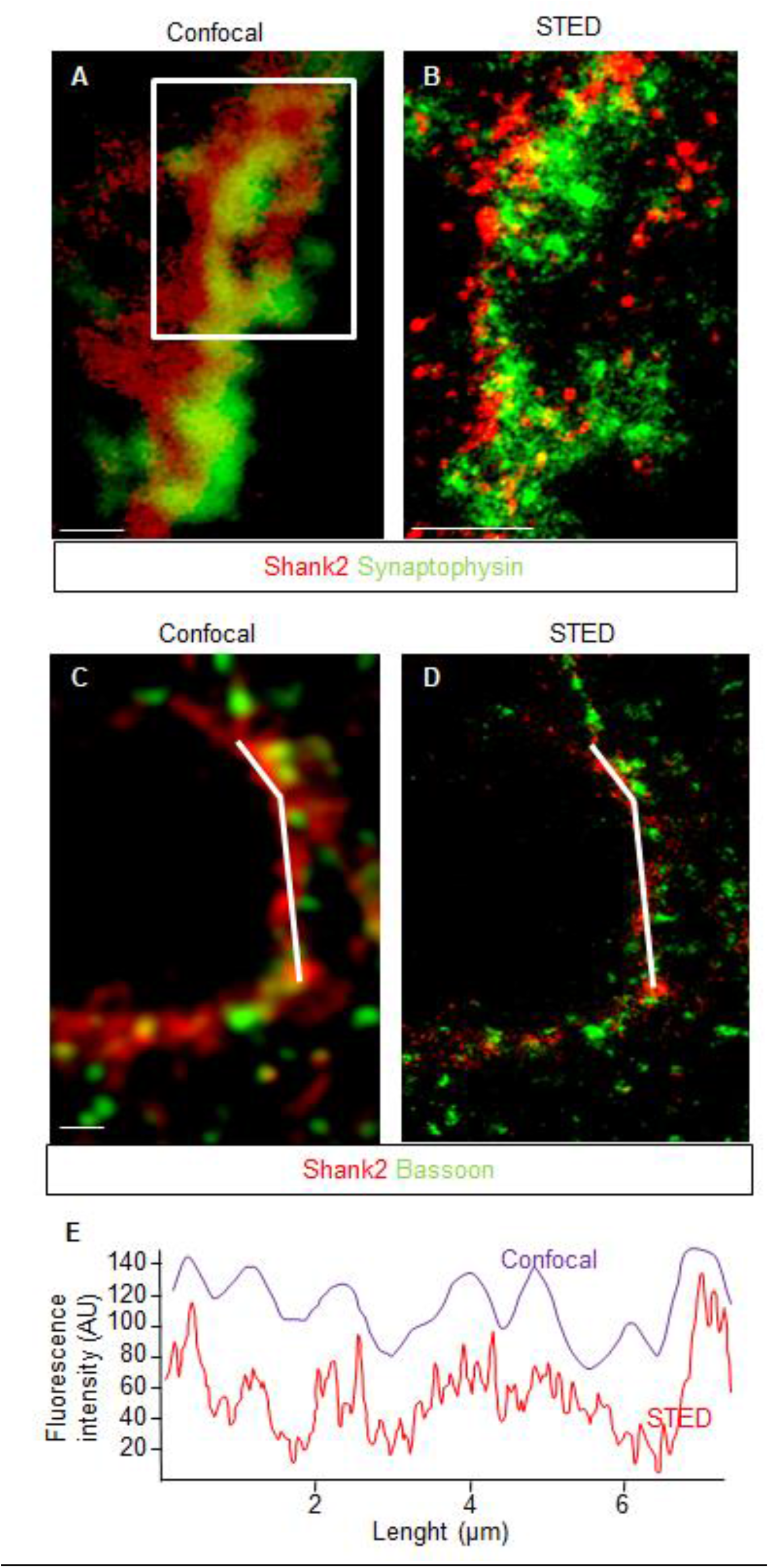
STED microscopy reveals punctuate structure of Shank2 colocalised with pre and post-synaptic structures. A) Confocal images of Shank2 and Synaptophysin show a band of Shank2 colocalized with Synaptophysin. B) STED images of Shank2 and Synaptophysin reveal the punctuate pattern of Shank2 and Synaptophysin. C) Confocal images of Shank2 and Bassoon show a band of Shank2 colocalized with Bassoon puncta. D) STED images of Shank2 and Bassoon reveal the punctuate pattern of Shank2 with Bassoon. Scale bar: A-B; 500nm, C-D; 1μm.

### Shank2+ synapses in Shank2^high^ neurons receive multiple mechanosensory and propriospinal input

We then set out to investigate if the Shank2 clusters on Shank2^high^ neurons constituted the specialized postsynaptic counterpart of a specific input, e.g. from dorsal root ganglia afferents (Li et al., 2011; Foster et al., 2015).

First, we co-immunostained spinal cord sections for Shank2 and for the presynaptic proteins vGluT1 (marker of cutaneous and mechanosensory projections) and vGluT2 (largely representing propriospinal projections; Alvarez et al., 2004) and we assessed if Shank2+ structures were preferentially juxtaposed to one of the two terminals. Interestingly, Shank2 immunopositivity in Shank2^high^ was not restricted to one of the two types of input: vGlut1+ presynaptic terminals covered 29.7±6.7% of Shank2+ cluster area, whereas vGlut2+ terminals covered no less than 65.6±8.2% (Figure 4A-C).

**Figure 4:**
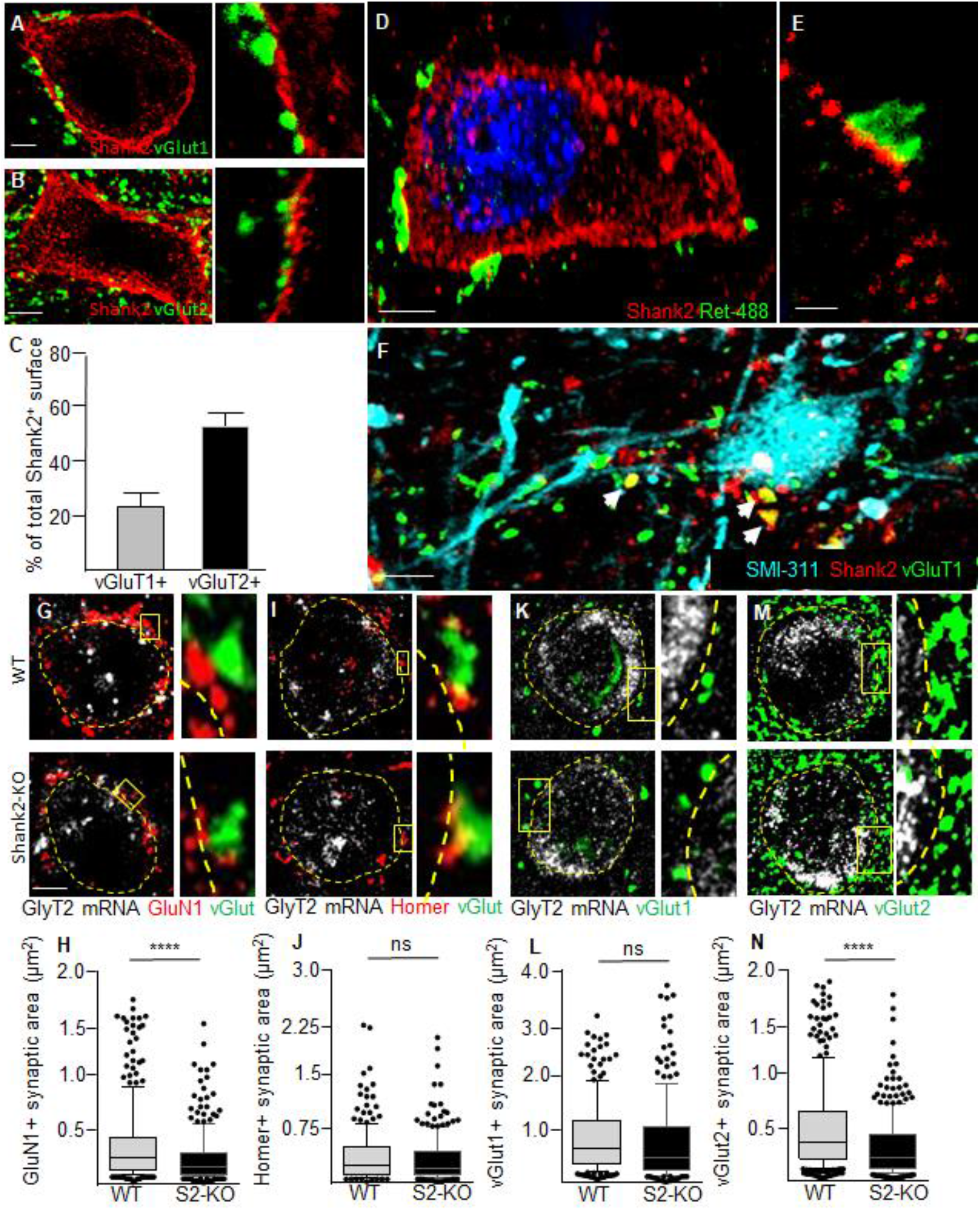
Propriospinal and mechanosensory input on Shank2^high^ expressing neurons. Synaptic input on Shank2 high expressing glycinergic neurons. A-C) Excitatory pre-synaptic input on Shank2 post-synaptic structures show both a vGlut1 (29.7±6.7% of total Shank2 area) and vGlut2 (65.6±8.2% of total Shank2 area) input on glycinergic interneurons. D-E) Detection of GFP positive buttons on Shank2 postive terminals in an c-RET-GFP mouse to show mechanosensory afferents on Shank2^high^ expressing neurons. F) IF staining on post mortem human spinal cord reveal that Shank2 post-synaps receive vGluT1 pre-synaptic input (36.9±14.1% of Shank2 positive puncta) G-J) post-synaptic structure GluN1 shows a significant decrease in size on glycinergic neurons in Shank2-KO mice (0.38±0.37μm^2^ vs 0.24±0.24μm^2^; WT vs KO; p<0.0001), however this decrease was not observed in the post-synaptic structure Homer (0.36±0.36μm^2^ vs 0.32±0.32μm^2^; WT vs KO; p>0.05). K-N) Pre-synaptic structure v-Glut1 was not affected on glycinergic cells in Shank2-KO mice (0.86±0.69μm^2^ vs 0.79±0.78μm^2^; WT vs KO; p>0.05), interestingly v-Glut2 shows a significant decrease in size on glycinergic neurons in Shank2-Ko mice (0.51±0.42 vs 0.32±0.28μm^2^; WT vs KO; p<0.0001). Data shown as average±SD, n=4, scale bar; A-B: 2,5μm, D: 5μm, E: 1μm, F: 10μm, G-M: 5μm.

We sought to more precisely confirm the nature of the projections to Shank2+ synapses. First, we injected fluorescently-labelled Cholera Toxin B subunit (CTB-488) into the hairy skin to label cutaneous afferents; central terminals labelled by this approach were distributed in a narrow column elongated along the dorso-ventral axis ranging from lamina II to lamina V (Li et al., 2011). Notably, even when a comparatively small area of the hindlimb skin was injected, several CTB-positive presynaptic boutons were detected on the cell body and proximal dendrites of Shank2^high^ glycinergic interneurons, in close apposition to Shank2 clusters (Supplementary Figure 2A-C).

Second, we used c-RET-GFP transgenic mouse (Hoshi et al., 2012) to track the central processes of mechanosensory afferents (Bourane et al., 2015). Also in this case, numerous GFP+ terminals were identified impinging on Shank2^high^ neurons; GFP+ presynaptic terminals were found to be juxtaposed to large Shank2 postsynaptic clusters (Figure 4D-E). Finally, immunostaining on post mortem human spinal cord revealed that the post-synaptic Shank2 terminals receive pre-synaptic input of vGluT1, 36.9±14.1% of Shank2 synapses were positive for vGlut1 (Figure 4F), which was comparable to the mouse data.

Thus, Shank2^high^ receive direct somatosensory input but Shank2+ synapses to not correspond to a distinct type of input.

**Supplementary Figure 2:**
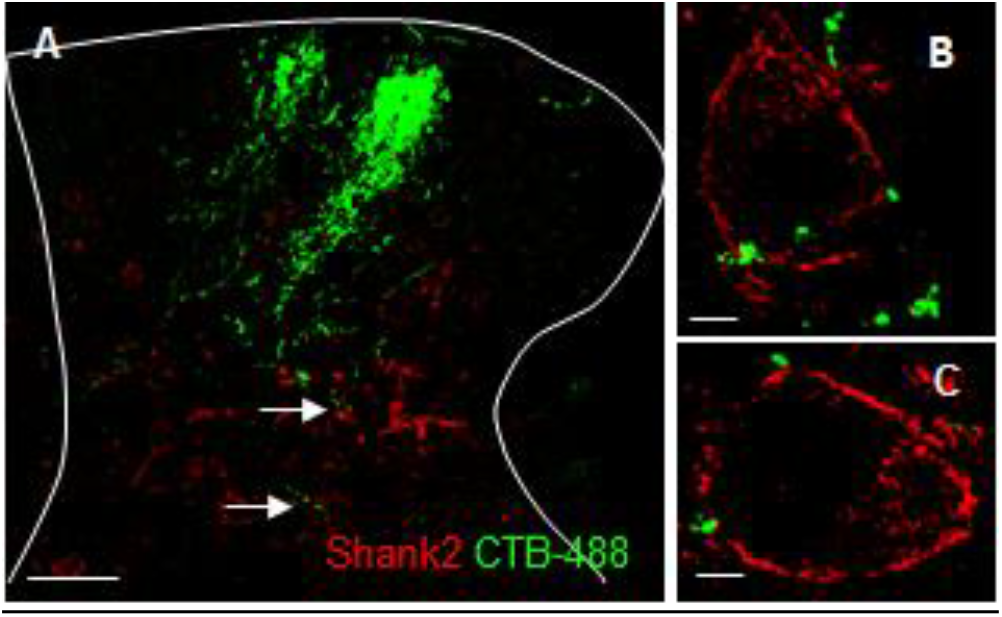
CTB-488 injections reveals cutaneous afferents on Shank2^high^ expresing neurons. A-C) Detection of GFP positive buttons on Shank2 positive terminals after Cholera Toxin B subunit injection in the hairy skin to show cutaneous afferents on Shank2^high^ expressing neurons. Scale bar; A: 50μm, B-C: 5μm.

### Neuronal architecture of the spinal cord is unaffected by Shank2-KO

Having established the existence and the nature of a new Shank2^high^ subpopulation of glycinergic interneurons, we set out to investigate how the loss of Shank2 in Shank2-KO mice may affect these interneurons and generate the pain-hypersensitivity phenotype observed in behavioural tests. Since several transgenic mouse lines with sensory or nociceptive phenotypes are characterized by disruption or selective loss of neuronal subpopulations (Ross et al., 2010, Wang et al., 2013) we verified that loss of Shank2 did not result in any gross disturbance of spinal cord architecture. In fact, the overall density of neurons (NeuN^+^; Supplementary Figure 3A-B) and in particular of inhibitory neurons (Pax2^+^) in the dorsal laminae of the spinal cord was comparable in WT and Shank2-KO (Supplementary Figure 3C-D). Likewise, the number of neurons expressing GlyT2 mRNA was similar in WT and Shank2-KO mice (Supplementary Figure 3G-H).

Finally, we determined the density of glycinergic synapses across the dorsal laminae, no significant difference has been found between WT and Shank2-KO (Supplementary Figure 4A-C). Thus, deletion of Shank2 did not cause any obvious changes in the overall architecture of the dorsal horn.

**Supplementary Figure 3:**
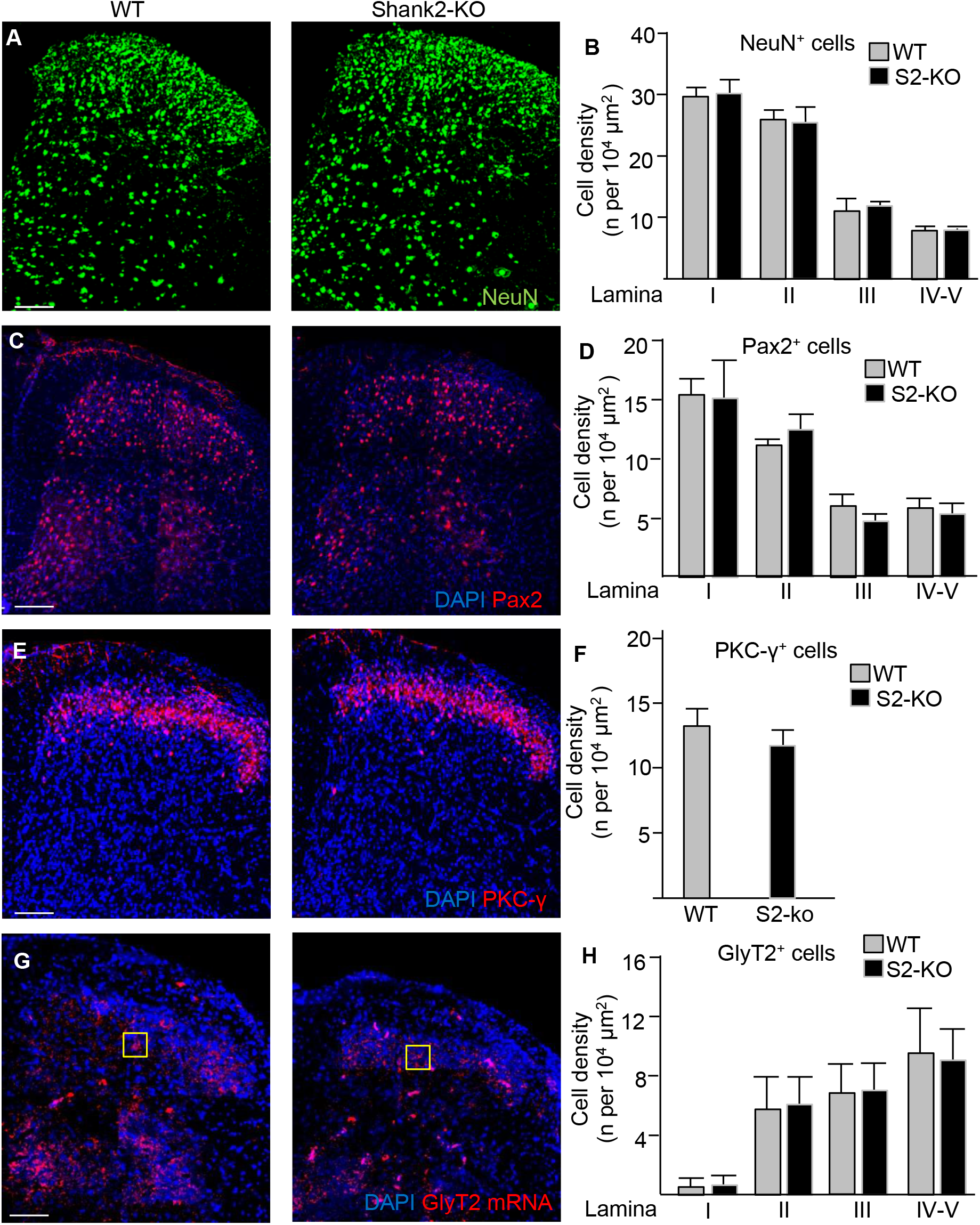
no difference in neuronal architecture in the spinal cord of Shank2-KO mice. A-B) Neu-N staining in the spinal cord reveals no differences in Neuronal counts between the WT and Shank2-KO across Laminea (29.4±1.9 vs 30.3±2.3; 26.1±1.6 vs 25.5±2.7; 11.1±2.1 vs 12.0±0.6; 7.8±0.8 vs 7.9±0.4; per 10^4^ μm^2^; Lamina I-IV/V respectively; WT vs KO; p>0.05). C-D) PAX2 staining in the spinal cord shows no differences in the density of inhibitory neurons between the WT and Shank2-KO (15.2±1.4 vs 15.0±3.2; 11.2±0.5 vs 12.2±0.7; 5.9±1.1 vs 4.7±0.7; 5.3±0.4 vs 5.7±1.0; per 10^4^ μm^2^; Lamina I-IV/V respectively; WT vs KO; p>0.05). E-F) PKC-y staining in the spinal cord results in no difference in the amount of excitatory cells between WT and Shank2-KO (13.2±1.5 vs 11.7±1.3; per 10^4^ μm^2^; WT vs KO; p>0.05). G-H) single mRNA in situ hybridisation with GlyT2 in the spinal cord reveals no differences in the density of Glycinergic interneurons between WT and Shank2-KO (0.1±0.1 vs 0.1±0.2; 1.4±0.5 vs 1.5±0.4; 1.6±0.5 vs 1.7±0.4; 2.2±0.7 vs 2.1±0.5; per 10^4^ μm^2^; Lamina I-IV/V respectively; WT vs KO; p>0.05). Data shown as average±SD, n=3, Scale bar: 100μm.

**Supplementary Figure 4:**
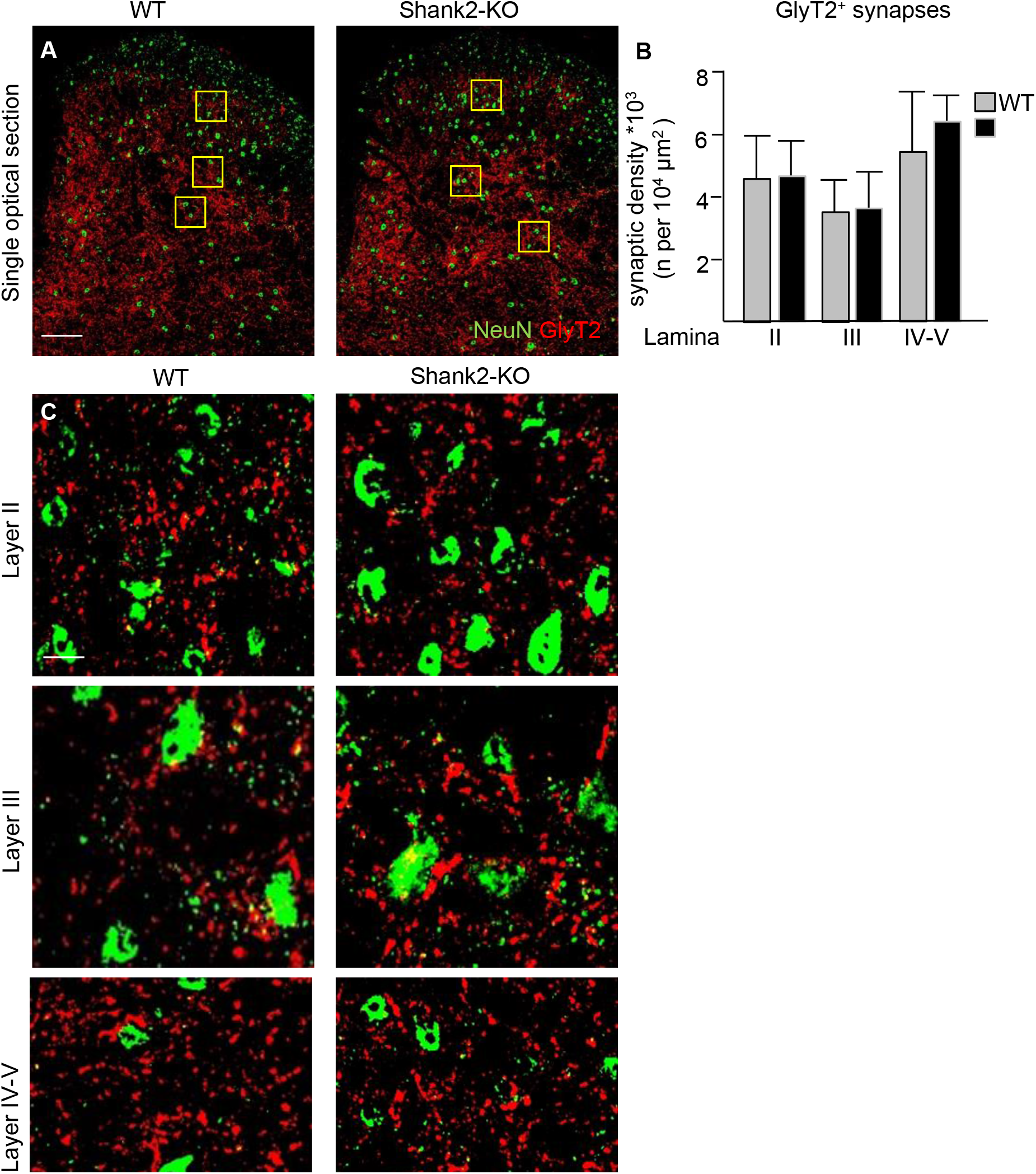
no difference in glycinergic synapses in the spinal cord of Shank2-KO mice. A-C)GlyT2 staining in the spinal cord reveals no differences in the GlyT2 synaptic density between WT and Shank2-KO across all lamina (45.1±15.7 vs 45.9±14.1; 35.3±11.6 vs 37.1±10.8; 54.6±20.8 vs 64.8±8.8;Lamina II, III and IV respectively; WT vs

### Loss of Shank2 disrupts NMDAR clustering in excitatory synapses on glycinergic interneurons

Since Shank2 is a core scaffold protein in glutamatergic synapses, we reasoned that excitatory synapses on Shank2^high^ glycinergic interneurons may suffer substantial postsynaptic (and/or presynaptic) alterations in Shank2-KO mice. We identified glycinergic interneurons by the detection of GlyT2 in in situ hybridization and co-immunostained the sections for post-synaptic proteins and receptors. Because of the protease treatment required for mRNA detection, only a restricted number of antibody-antigen couples (whose antigens were protease-resistant) could be employed. In particular, we considered the GluN1 NMDAR subunit and the PSD scaffold protein Homer. Interestingly, the number of GluN1 and Homer post-synaptic structures on GlyT2 interneurons was comparable in WT vs Shank2-KO mice (95.3±17.5 vs 107.4±31 per 100μm WT vs KO for GluN1; 80.1±20.2 vs 104.7±25.8 per 100μm WT vs KO for Homer) indicating that the number of synapses and their qualitative composition were not affected by Shank2 loss. However, GluN1 clusters were significantly smaller in glycinergic interneurons of Shank2-KO mice compared to WT mice (p<0.0001, Figure 4G-H). Remarkably, this reduction in size was not observed for Homer (p>0.05; Figure 4I-J)

Since Shank2 proteins may regulate trans-synaptic signaling, we also took into consideration the number and the size of presynaptic excitatory boutons. To this aim, glycinergic interneurons were identified by GlyT2 in situ hybridization and vGlut1+ and vGlut2+ terminals by immunostaining. The number of vGlut1 and vGlut2 terminals on GlyT2 interneurons was comparable in WT and Shank2-KO mice (16.3±5.2 vs 17.8±7.8 per 100μm WT vs KO for vGlut1; 59.2±17.7 vs 64.2±18.8 per 100μm WT vs KO for vGlut2); however, vGluT2 terminals were significantly smaller in Shank2-KO mice compared to WT mice (p<0.0001; Figure 4M-N). This difference was not observed in the vGluT1 terminals (p>0.05; Figure 4K-L).

Taken together these data show that loss of Shank2 does not affect the number of excitatory synapses on glycinergic interneurons but causes the selective decrease of synaptic NMDAR content and in the size of propriospinal glutamatergic presynaptic terminals. Thus, Shank2 loss appears to selectively weaken glutamatergic input and plasticity on glycinergic interneurons.

### Reduced activation of glycinergic interneurons in Shank2-KO mice is associated with increased excitation of dorsal laminae interneurons upon nociceptive stimulation

Next we investigated if the disturbance in the structure of synaptic inputs to Shank2^high^ interneurons affected their activation during nociceptive stimulation. To this aim, WT and Shank2-KO mice were injected with Formalin (or with saline) in the plantar aspect of the right hindpaw (as in the Formalin test) and sacrificed 120 min later (90 min after the onset of the phase-II). We used single-mRNA molecule in situ detection to simultaneously identify glycinergic interneurons (using GlyT2 mRNA) and c-fos mRNA expression as a marker of neuronal activity (the use of c-fos immunostaining was precluded by the protease treatment necessary for the identification of GlyT2 mRNA). Indeed, the number of c-fos^+^/GlyT2^+^ neurons was very low at baseline and comparable in WT and Shank2-KO samples (p>0.05 Figure 5A-B); however, upon Formalin challenge, the number of double-positive interneurons increased strongly in WT animals but much less so in Shank2-KO (p<0.01; Figure 5A-B) implying a failure in GlyT2+ neuron activation upon nociceptive stimulation in the Shank2-KO mice.

**Figure 5:**
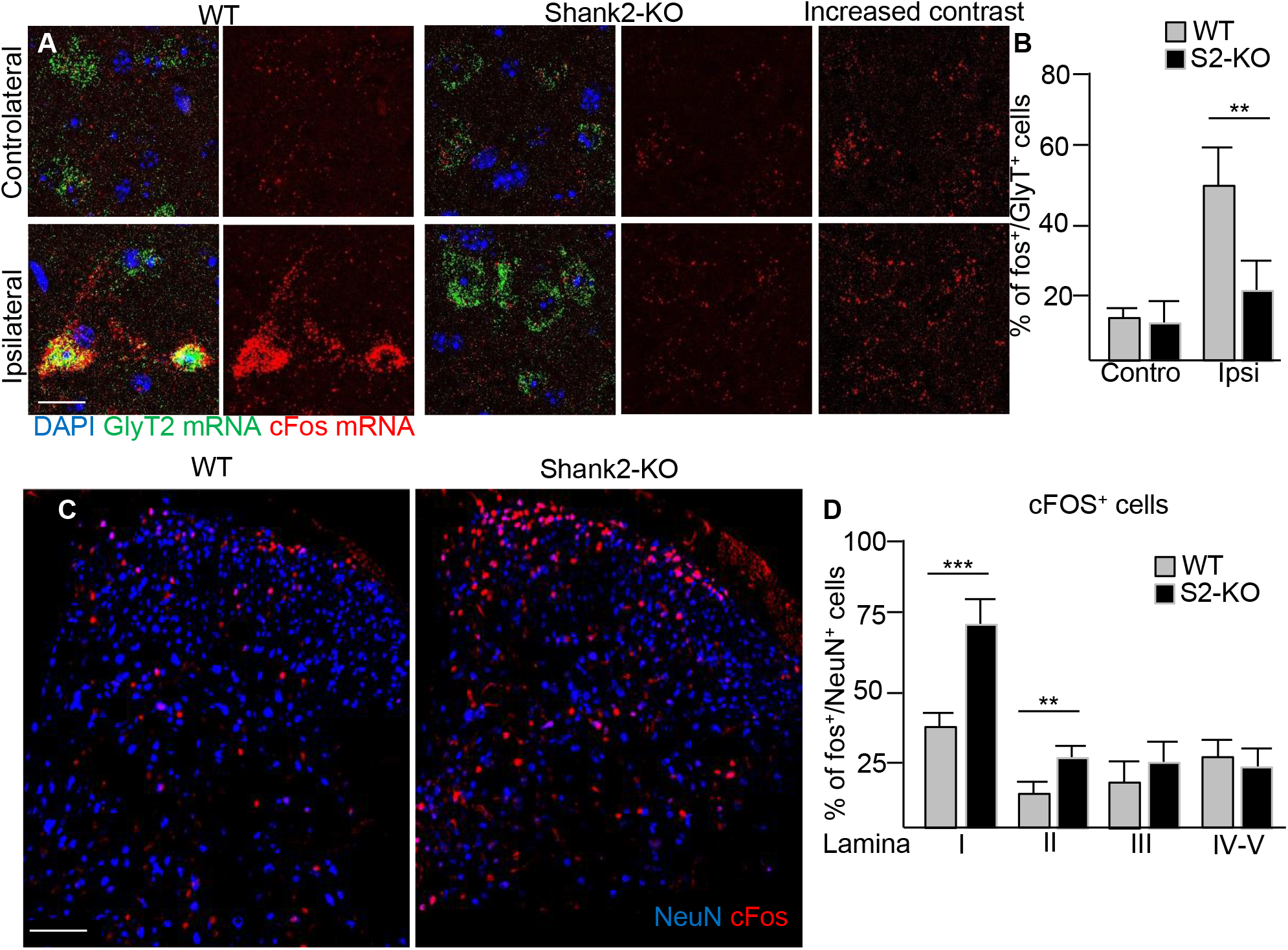
Reduced activity in glycinergic interneurons is associated with increased activity of interneurons in the dorsal lamina I and II. Activation patterns in glycinergic interneurons and interneurons in dorsal lamina upon activation. Mice were treated with low concentration of Formalin and c-fos levels were measured 120min later A-B) Saline injection resulted in no difference in c-fos levels in glycinergic cells between WT and KO mice (10.7±4.7% vs 9.8±5.2%; WT vs KO; p>0.05). Upon Formalin injection c-fos was significantly higher in WT vs KO mice in glycinergic cells (50.6±7.3% vs 18.1±6.6%; WT vs KO; p<0.01). C-D) The reduced activity of glycinergic cells in Shank2-KO mice in turn significantly increased the overall c-fos expression in the KO in lamina I (33.5±8.5% vs 71.5±10.5%; WT vs KO; p<0.001), Lamina II (13.5±2.4% vs 23.2±7.5%; WT vs KO; p<0.01) and a strong trend in layer III (15.6±7.6% vs 25.5±12.2%; WT vs KO; p>0.05) upon formalin injection. Data shown as average±SD, n=4, scale bar; A: 20μm, C: 100μm.

We then sought to verify if the observed decrease in inhibitory interneurons activity would result in the overall increase in excitation in the dorsal-horn circuitry of Shank2-KO mice. We considered spinal cord sections obtained 120 min after a Formalin challenge in the right hindpaw; in order to monitor the large-scale activation of neurons in the dorsal spinal cord, the samples immunostained for NeuN (to identify all neurons) and c-Fos and the ratio of c-Fos^+^/NeuN^+^ cells was annotated according to the spinal cord lamina. In saline-injected WT mice, c-Fos^+^ neurons were detectable in very low numbers across dorsal spinal cord (2.2±1.1% in lamina I) and in saline-injected Shank2-KO mice the percentage of c-fos^+^ neurons was comparable to WT mice (4.5±2.7 % of NeuN^+^ in lamina I; p>0.05). Formalin injection caused a much larger elevation of c-Fos^+^ neurons in Shank2-KO vs WT in lamina I (p<0.001; Figure 5C-D) as well as in lamina II (p<0.01, Figure 5C-D); because of the high variability of the size of the c-Fos^+^ population, a strong trend was detected for lamina III (p>0.05; Figure 5C-D) but no difference was found in lamina IV and V (Figure 5C-D). A comparable result was obtained when the number of cells positive for c-Fos mRNA (by in situ hybridization) was measured in the same groups, underscoring the robustness of the dataset (Supplementary Figure 5A-B).

Taken together, these data suggest that the disruption of the excitation of inhibitory interneurons in Shank2-KO mice results in a decreased excitation of glycinergic inhibitory interneurons upon acute chemical pain and, in turn, to the increased activation of neurons in the dorsal horn following a nociceptive stimulus.

**Supplementary Figure 5:**
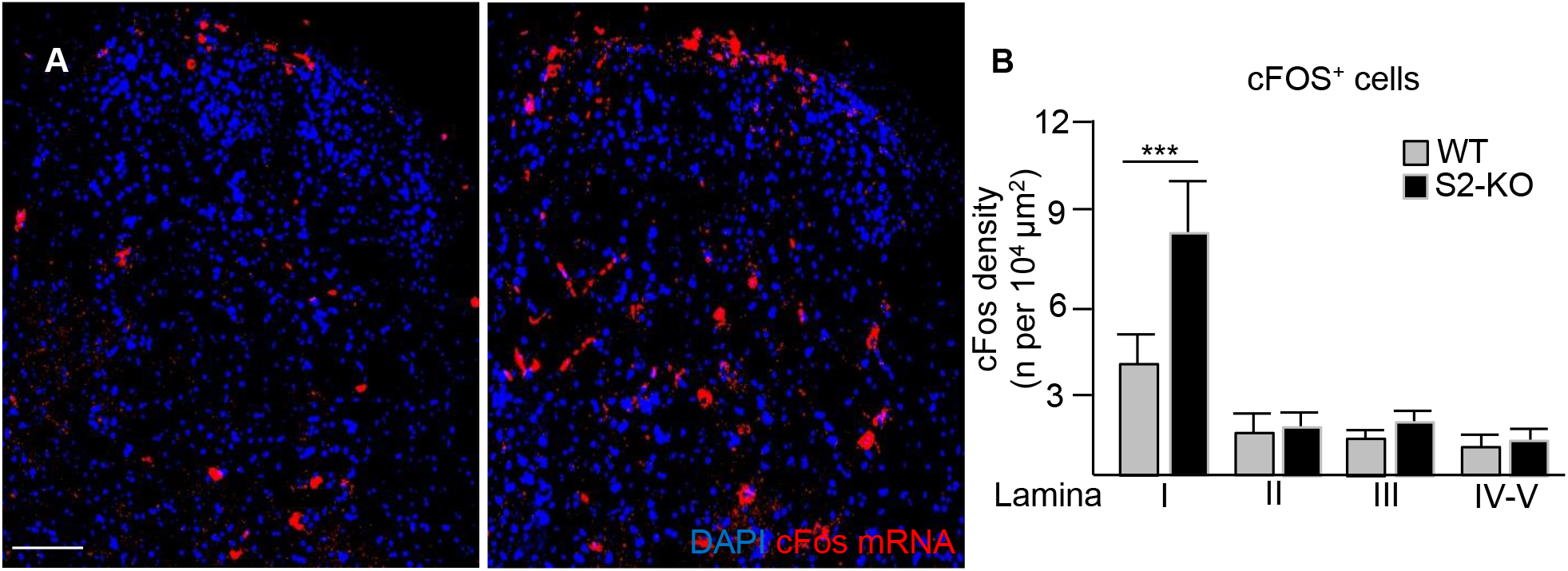
Increased number of cFos mRNA+ interneurons in Lamina I upon Formalin injection. A-B) Single mRNA detection show a strong increase in c-fos expressing cells in lamina I (4.0±0.9 vs 8.7±1.4; WT vs KO; p<0.001), however this effect dissapears in Lamina II-IV/V (1.5±0.6 vs 1.6±0.5; 1.2±0.3 vs 1.7±0.4; 0.8±0.2 vs 1.1±0,3; per 10^4^ μm^2^; Lamina II-IV/V respectively; WT vs KO; p>0.05) upon formalin injection. Data shown as average±SD, n=4, scale bar; 100μm.

## Discussion

Our data show that high levels of Shank2 expression identifies a subpopulation of glycinergic inhibitory interneurons located in the dorsal horn of the spinal cord, receiving proprioceptive inputs from somatosensory afferents; Shank2 loss leads to a decrease in glycinergic interneurons excitatory synapses and in net increase of excitation in the dorsal horn, correlated with increased sensitivity to chemically-induced pain.

Genetic approaches have increasingly revealed the high degree of heterogeneity in neuronal subpopulations in the dorsal spinal cord (Ross et al., 2010; Wang et al., 2013; Bourane et al., 2015). Recently, the genetic diversity of these populations has been demonstrated to be extensive and a large number of subpopulations have been identified in a single-cell transcriptome study (Häring et al., 2018). Nevertheless, neuronal physiology is highly influenced by the quantity and quality of their synaptic inputs, which is strongly dependent upon the composition and architectural organization of post-synaptic structures (Schlüter et al., 2006; Favaro et al., 2018). Therefore, a distinct layer of diversity may be identified once the synaptic composition of neuronal subtypes is taken into consideration. Here we provide a first proof of this concept; in fact, Shank2 distinguishes a subclass of glycinergic and parvalbumin interneurons as well as a small population of excitatory interneurons.

Since different members of the Shank family, despite their similarity, are not considered mutually redundant (Shi et al., 2017; Schmeisser et al., 2013), the function of Shank2^high^ cells is predicted to be heavily impacted by Shank2 loss, despite their anatomical integrity (i.e., their normal number and positioning). In agreement with observations in other neuronal subtypes (Chung et al., 2019; Shi et al., 2017; Pappas et al., 2017), we find that loss of Shank2 causes the decrease in NMDAR expression in excitatory synapses on GlyT2+ interneurons. Although baseline neurotransmission in the pain processing circuit appears to be not affected (as the acute phase of the formalin test is comparable in Shank2-KO mice and WT littermates), the NMDAR-dependent synaptic plasticity that is thought to underlie the second phase of the behavioural response to the formalin test (Coderre et al., 1990; Ji et al., 1999; Asante et al., 2009) may be unbalanced, with insufficient potentiation of the inhibitory circuit. In fact, glycinergic interneurons are strongly activated in WT (as shown by the c-Fos induction), but not in the Shank2-KO animals. Thus, the resulting decrease in glycinergic transmission in the pain processing circuit would cause excessive excitatory drive, as demonstrated by the increase in the number of c-fos+ neurons in lamina I and II. In fact, dysfunction or silencing of inhibitory interneurons is known to cause pain hypersensitivity in human patients and experimental animals (Vuillemier et al., 2018; Imlach et al., 2016; Foster et al., 2015) in particular by dis-inhibiting PKC-gamma+ excitatory interneurons (Lu et al., 2013).

The nociceptive disturbance of Shank2-KO animal models is actually multidimensional. In fact, Shank2-KO mice have been reported to display a reduced allodynia response (higher mechanical threshold for withdraw 3 and 7 days after Sciatic Nerve Ligation -SNL- or after injection of Complete Freund Adjuvant-CFA) upon induction of chronic inflammatory or neuropathic pain, whereas at baseline a reduced sensitivity to heat-induced pain was detected (Ko et al., 2016). Our findings are in agreement with the reported reduced sensitivity to allodynia induction upon CFA injection (Figure 1D) and extend the characterization of the phenotype showing a hypersensitivity to pain in the formalin test and in heat-sensitivity upon formalin injection. Thus, depending on the acute or chronic phase and on the type of noxious stimulus, loss of Shank2 may result in increased or decreased sensitivity. Interestingly, Shank2-KO mice have been also reported to display a reduced sensitivity to the nociceptive response evoked by intrathecal injection of NMDA (Yoon et al., 2017). Since this procedure does not selectively activate one subpopulation of neurons, it is not possible to explain it in circuit terms. In fact, Shank2 is highly enriched in glycinergic interneurons, but it is not restricted to these cells (see Figure 1A-B) and even among the Shank2^high^ cells, a small fraction of excitatory neurons (VGluT2^+^, Prrxl1^+^) can be identified and their role remains unexplored. Furthermore, the dysfunction of PV interneurons in spinal cord has been directly related to mechanical but not thermal allodynia (Petitjean et al., 2015); interestingly, only a fraction of PV interneurons appear to be Shank2^high^ and indeed we detect thermal but not mechanical allodynia.

Thus, the impact of Shank2 loss may affect modality-specific pain processing circuits in a distinct way, depending on the role of different cellular subpopulations.

The insufficient activation of glycinergic interneurons because of disrupted excitatory synapses observed in Shank2-KO mice is a new mechanism for abnormal pain processing in autism. In fact, reported pain hyposensitivity in Shank3 mice has been linked to the loss of Shank3 in neurons in the dorsal root ganglia and the direct effect of Shank3 absence on TRPV channel expression (Han et al., 2016). Likewise, autism-related behavioural dysfunctions have been linked to the disturbed sensory input generated by abnormal sensory neurons in dorsal root ganglia (Orefice et al., 2016; Orefice et al., 2019). Conversely, loss of function of the ASD-associated gene Caspr2 is associated with neuropathic pain (Dawes et al., 2018) through mechanisms involving increased sensitization of neurons in dorsal root ganglia. Although these reports all point to a sensory dysfunction originating in periphery, our findings suggest that disruption of spinal cord circuits may be a strong contributor to the observed hyper- and hyposensitivity to nociceptive stimuli. Furthermore, one can speculate that the same excitation/inhibition balance disruption that is thought to underlie the ASD spectrum disorder may also manifest itself in spinal circuits to contribute to drive sensory abnormalities.

In conclusion, we demonstrate that the Shank2 expression level characterizes a subset of inhibitory interneurons whose activation is disrupted by Shank2 loss leading to excessive nociceptive circuit activation and allodynia. Although Shank3 expression is known to be compensatorily increased upon Shank2 KO (Schmeisser et al., 2012), no functional compensation takes places in glycinergic interneurons, whose excitatory synapses appear to be functionally and structurally disrupted. It remains unclear which role of Shank2 is absolutely required in these cells, that cannot be compensated by other Shank family members. Our findings also suggest that treatment of nociceptive disturbances in ASD may require to be tailored to the underlying genetic cause, even when phenotypes converge, since distinctive pathophysiology may be at play in each case.

## Acknowledgement

FR and TB are supported by the DFG in the context of the CRC1149 “Danger Response, Disturbance Factors and Regenerative Potential after Acute Trauma”. FR is also supported by the Baustein program of Ulm University, the Synapsis Foundation, the Radala foundation and the Thierry Latran Foundation. The authors would like to thank prof. Frank Kirchoff for the use of the confocal facility and prof. Anita Ignatius for the access to the histology facility. The technical assistance provided by Thomas Lenk is gratefully acknowledged.

## Notes

### Competing Interest Statement

The authors have declared no competing interest.

